# Accounting for GC-content bias reduces systematic errors and batch effects in ChIP-Seq data

**DOI:** 10.1101/090704

**Authors:** Mingxiang Teng, Rafael A. Irizarry

## Abstract

The main application of ChIP-seq technology is the detection of genomic regions that bind to a protein of interest. A large part of functional genomics public catalogs are based on ChIP-seq data. These catalogs rely on peak calling algorithms that infer protein-binding sites by detecting genomic regions associated with more mapped reads (coverage) than expected by chance as a result of the experimental protocol's lack of perfect specificity. We find that GC-content bias accounts for substantial variability in the observed coverage for ChIP-Seq experiments and that this variability leads to false-positive peak calls. More concerning is that GC-effect varies across experiments, with the effect strong enough to result in a substantial number of peaks called differently when different laboratories perform experiments on the same cell-line. However, accounting for GC-content in ChIP-Seq is challenging because the binding sites of interest tend to be more common in high GC-content regions, which confounds real biological signal with the unwanted variability. To account for this challenge we introduce a statistical approach that accounts for GC-effects on both non-specific noise and signal induced by the binding site. The method can be used to account for this bias in binding quantification as well to improve existing peak calling algorithms. We use this approach to show a reduction in false positive peaks as well as improved consistency across laboratories.

## Introduction

Chromatin immunoprecipitation followed by NGS (ChIP-Seq) is widely used for detecting the genomic locations of transcription factor binding and histone modifications. ChIP-Seq is widely used, with the majority of data provided by the ENCODE (Dunham et al. 2012) and modENCODE (Celniker et al. 2009) projects produced with this technology. After mapping the NGS reads, the main part of the quantitative analysis is to infer the genomic sites where the protein of interest binds by finding regions with an enrichment of mapped reads. The regions reported by this analysis are referred to as peaks due to the appearance of the coverage plots (Pepke et al. 2009). Several competing peak detection algorithms have been described in the literature (Ji et al. 2008; Jothi et al. 2008; Kharchenko et al. 2008; Valouev et al. 2008; Zhang et al. 2008; Rozowsky et al. 2009; Rashid et al. 2011). Although details of these competing approaches vary, most follow similar general principles. First, after reads are mapped, coverage is calculated for binned regions of the genome. In principle only regions including binding sites should have counts larger than 0. But due to non-specificity of the experimental protocol we observe a *background level*. This background level is then modeled and statistical inference is used to distinguish between count levels that can be explained with the background model and those that are higher than expected by chance. The latter are reported as peaks.

GC content bias has been reported for several NGS applications (Dohm et al. 2008; Alkan et al. 2009; Cheung et al. 2011; Benjamini and Speed 2012). For genomic DNA data, PCR amplification of DNA fragments during library preparation is one factor that introduces this bias (Aird et al. 2011; Ross et al. 2013). The bias has also been observed in RNA-Seq data (Love et al. 2016). Solutions to this bias have been published for genomic DNA (Benjamini and Speed 2012) and RNA-Seq data (Hansen et al. 2012; Love et al. 2016). However, below we explain why these approaches are not directly applicable to ChIP-Seq data.

Using ENCODE (Dunham et al. 2012) data we show that GC-content bias is also present in ChIP-Seq technology. Furthermore, we demonstrate that the way in which GC-content affects coverage varies across samples and laboratories and that this unwanted variability is substantial enough to results in different laboratories calling different regions as peaks. Unfortunately, solutions for GC-bias correction, published for other NGS applications, are not directly applicable to ChIP-Seq experiments. This is because, in many instances, binding sites are expected to occur in or near high GC-content regions such as gene promoters. If we naively correct for GC-content we may erase the biologically relevant signals we are interested in detecting. Here we present an approach, based on a mixture model, that accounts for GC-content bias separately for effects related to protein binding and differential non-specific binding. We demonstrate how this approach greatly reduces false positive peaks and improves agreement across laboratories.

## Results

### GC affects coverage and it does so differently in different labs

To demonstrate the challenges presented by GC-content bias and the advantages presented by our method, we downloaded and processed raw ENCODE (Dunham et al. 2012) ChIP-Seq data measuring CTCF binding for the GM12878, HeLa-S3, HepG2, HUVEC, K562 and NHEK cell lines. We selected this particular example because data is available from experiments performed by three independent laboratories each running two replicated experiments with at least one million mapped reads (cell lines GM12878 and K562 have 3 replicates from one lab).

To explore the extent and characteristics of the bias, we segmented the genome into 10K basepair bins (we used a large bin size because it facilitated exploratory data analysis) and GC-content was computed for each bin. Then for each of the HUVEC cell lines, we counted read starts for each bin. Plotting counts versus GC-content reveals two clusters in each of the samples (Fig. 1A-F). We used the HUVEC data to illustrate the challenge because it exhibited the strongest GC-content bias. The presence of two clusters are in agreement with the previously noted observation that ChIP-Seq reads can result from either 1) a background level or 2) protein binding regions (Zhang et al. 2008), with the latter associated with peaks. In both replicates from one laboratory we observe that counts increase with GC-content in both background and signal clusters (Fig. 1A and 1B). Of particular concern is the fact that the way GC-content affects coverage is different in another laboratory (Fig. 1C and 1D) in which counts decrease with GC-content. In a third laboratory, the GC-content bias is only present in one of the two replicates (Fig. 1E and 1F).

**Figure 1.**
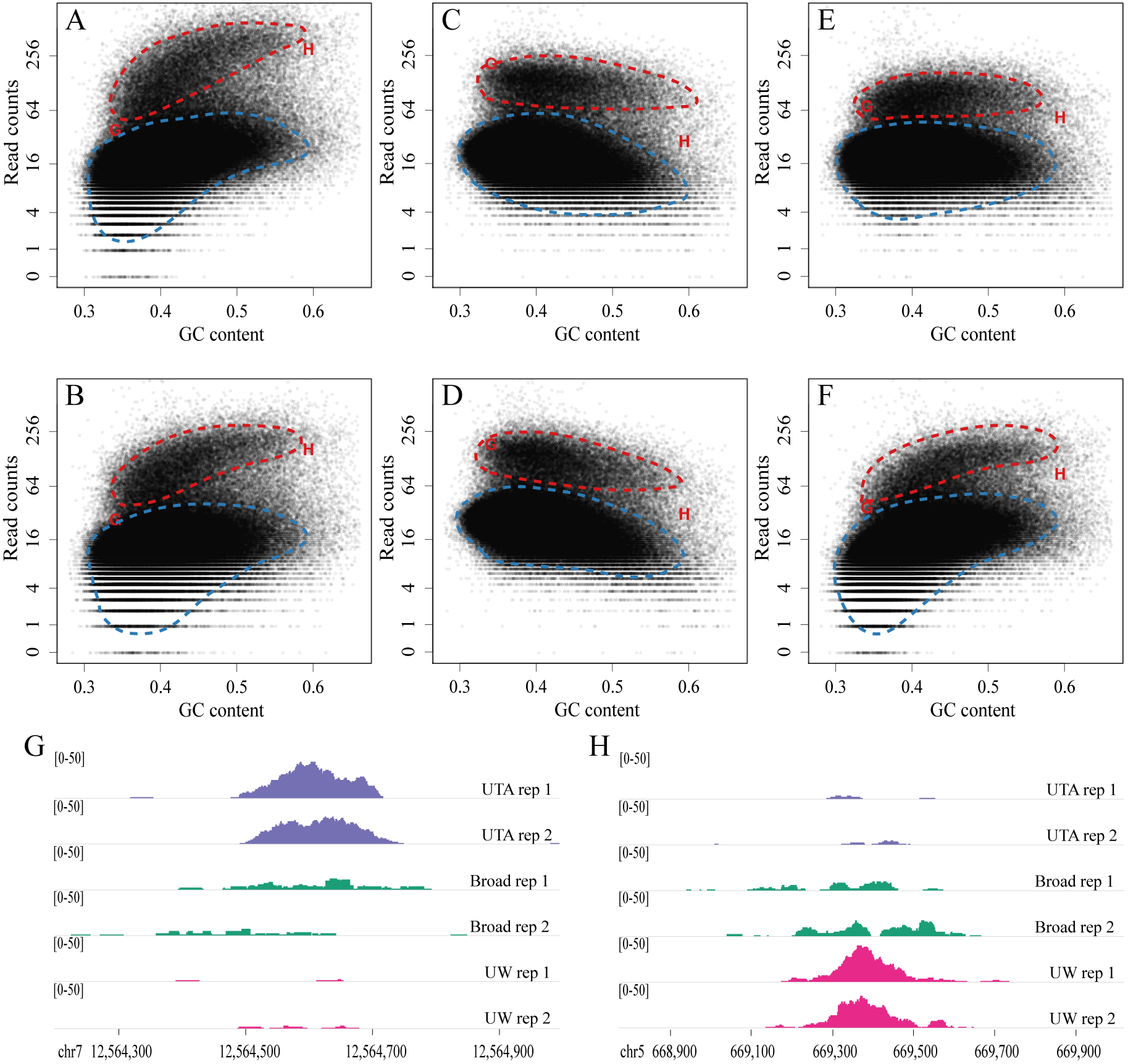
Evidence of GC-content effects at the bin level and its downstream result on peaks demonstrated on the HUVEC cell line. A) The genome is divided into 10 kB bins and counts are computed in the first replicate of laboratory UW each as well as the GC-content of each bin. Counts are plotted against GC-content. Hand-drawn contours are added to highlight the presence of two clusters. B) As A) but for the second replicate. C) as A) but for the first replicate of laboratory UTA. D) As C) but for the second replicate. E) as A) but for the first replicate of laboratory Broad. F) As E) but for the second replicate. G) An example of peak that changes substantially from laboratory to laboratory. This peak is associated with the bin denoted with an ‘G’ in A-F. H) As G but for the bin denoted with an ‘H’ in A-F.

### GC bias leads to variability of ChIP-Seq peak calling

These effects described above are strong enough to affect downstream analysis such as peak detection. For example, coverage can change drastically across laboratories depending on the GC-content of the region (Fig. 1G and 1H, Supplemental Fig. 1). Note that in the high GC-content region, laboratory UW shows a peak but laboratory UTA does not, while in the low GC-content region it is the other way around. These regions are not isolated examples. In fact, the agreement in peak calls across laboratories (Supplemental Fig. 2A) is rather low. For example, the peaks reported for the HUVEC cell-line on the ENCODE(Landt et al. 2012) portal report in 37,920, 44,033 and 37,412 peaks called for the three laboratories respectively with 24.3% of regions reported by only one lab. To see that GC-content was a major driver of these differences we compared the GC-content of the peak regions detected just by laboratory UW, to those detected just by laboratory Broad, to those detected just by laboratory UTA and found a strong difference (Fig. 2A). Note that these differences cannot be due to biology but rather must be a result of differences in experimental conditions. These results demonstrate that, if left unaccounted for, GC-content bias will lead current peak callers to report a substantial number of false positives.

**Figure 2.**
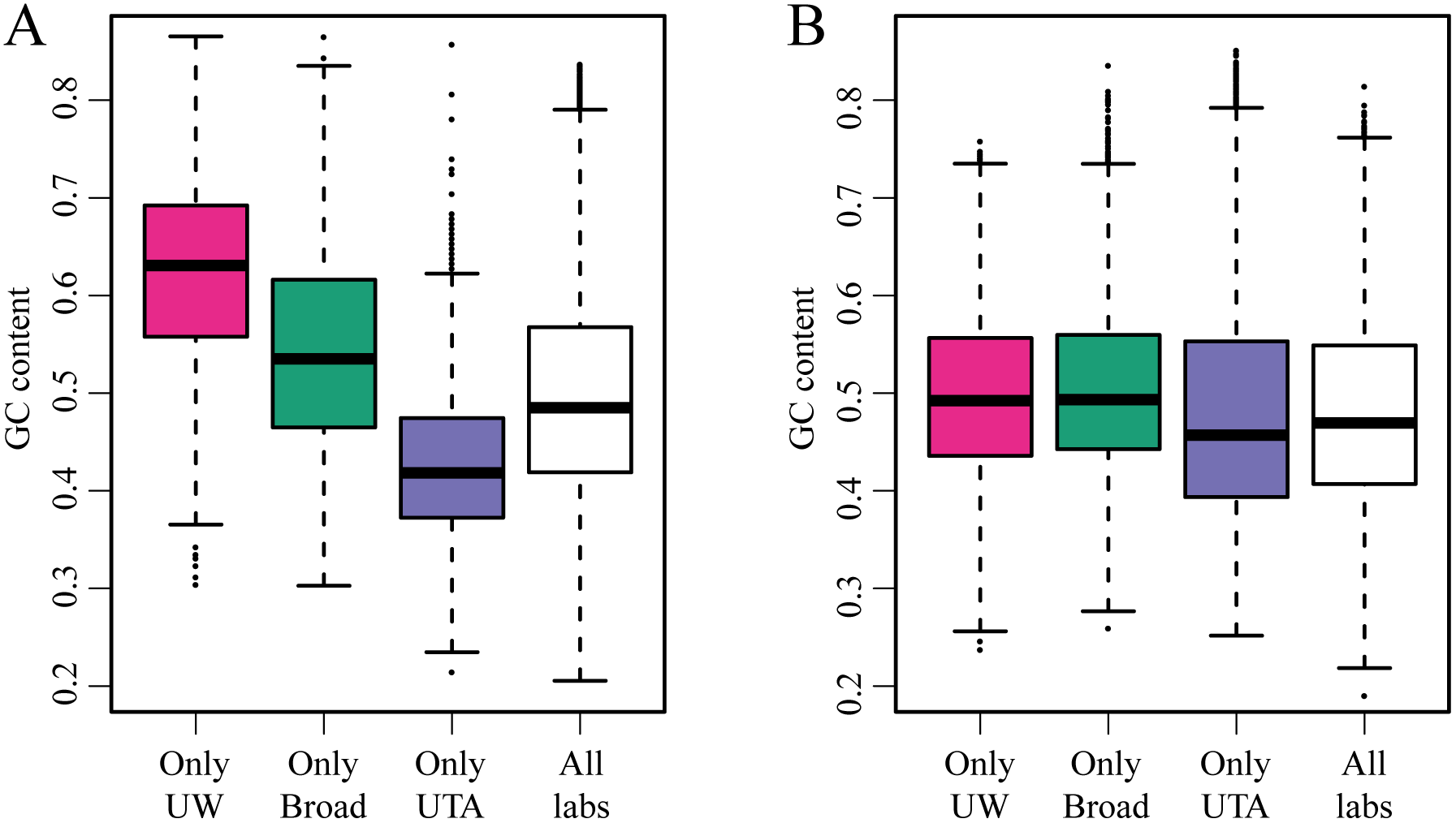
GC-content of peaks called by only one laboratory. (A) For the HUVEC cell line we formed four groups of peaks reported by ENCODE portal. We split them into those called only in laboratory UW, those called only in laboratory Broad, those called only in laboratory UTA and those called in all three. We computed the GC-content for each of these peak regions and show in four boxplots. (B) As (A) but after finding peaks with our algorithm.

### Mixture Model Estimates GC-content effect for background and signal

Published work on GC-content bias correction has found that modeling GC-content effects at the fragment level is, currently, the optimal approach (Benjamini and Speed 2012; Love et al. 2016). However, this approach is not directly applicable to ChIP-Seq data. One reason is that most peak calling algorithms operate on bin level information. Specifically, these algorithms define bins, compute coverage in these bins, and then peaks are inferred from these coverage measurements (Ji et al. 2008; Kharchenko et al. 2008; Zhang et al. 2008; Rashid et al. 2011). Here we develop a method that makes use of an approximation that permits the adaptation of published peak calling algorithms so that they adjust for GC-content bias. Although we focus on the SPP (algorithm) (Kharchenko et al. 2008) because it was used by the ENCODE project, our approach applies to any peak algorithm based on coverage computed in bins. The approach can also be used to adjust enrichment scores for predefined regions.

The first step of our approach is to estimate a sample specific GC-content effect from the data. This effect is defined by estimating the GC-bias for both background level and binding signal for any given potential binding region (Fig. 3). Suppose our targeted protein binds to a region centered at genomic location *i*_0_ and has length *l*. Computing the GC-content of the genomic region starting at 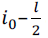 and ending at 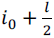 is straightforward. However, due to the fact that DNA is randomly cut into fragments of sizes ranging from 200bp to 500bp (depending on the experimental protocol) the sequenced reads associated with this binding site maps to a larger region of the genome (Fig. 3). Specifically, if fragments are, on average, size *h* then the peak region will span from 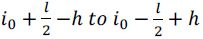. Note also that once outside of the 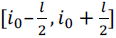 range, the probability that a specific fragments appears decreases as its center is further to *i*_0_. Specifically, these different probabilities imply that we should use the following weights:

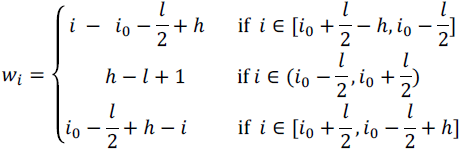

The shape of *w_i_* can be seen in Supplemental Figure 3. This implies that the total GC-content bias affecting fragments associated with a protein binding at 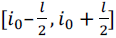 is a weighted average of all the GC-content effects of all potential fragments in the bin 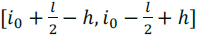. We therefore define the *effective GC-content* (EGCC) associated with a bin centered at *i*_0_ as

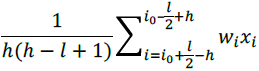

where *x_i_* is 1 if genomic location *i* is G or C, and 0 otherwise. Note that positive side effect of this approach is that it results in a GC-content covariate that is less sensitive to the bin size (Supplemental Fig. 4).

**Figure 3.**
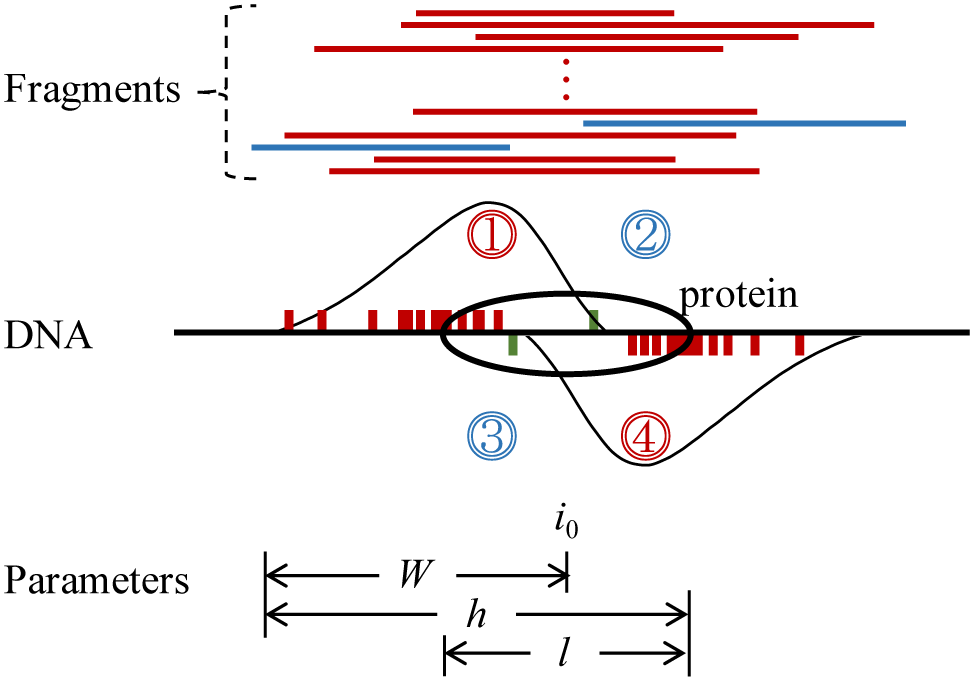
Illustration of regions related to the enrichment score calculation. The regions associated with the counts denoted by *Y_i, +_*, *Y_i, −_*, *B_i, +_*, *B_t, −_* in the paper are denoted with the regions 1, 4, 2, and 3 respectively.

With an EGCC in place for any given genomic location, we then estimate GC-content effects for both the background level and signal using a mixture model. Specifically, we pose a mixed generalized linear model with two components corresponding to coverage due to specific binding and background regions respectively. We assume that each component follows a Poisson distribution with the log of the rate a smooth function of EGCC. When fitting this model to the HUVEC data, the fitted GC-content dependent effects demonstrate that each laboratory introduces a different type of bias for both the signal and background (Fig. 4). See Methods Section for details.

**Figure 4.**
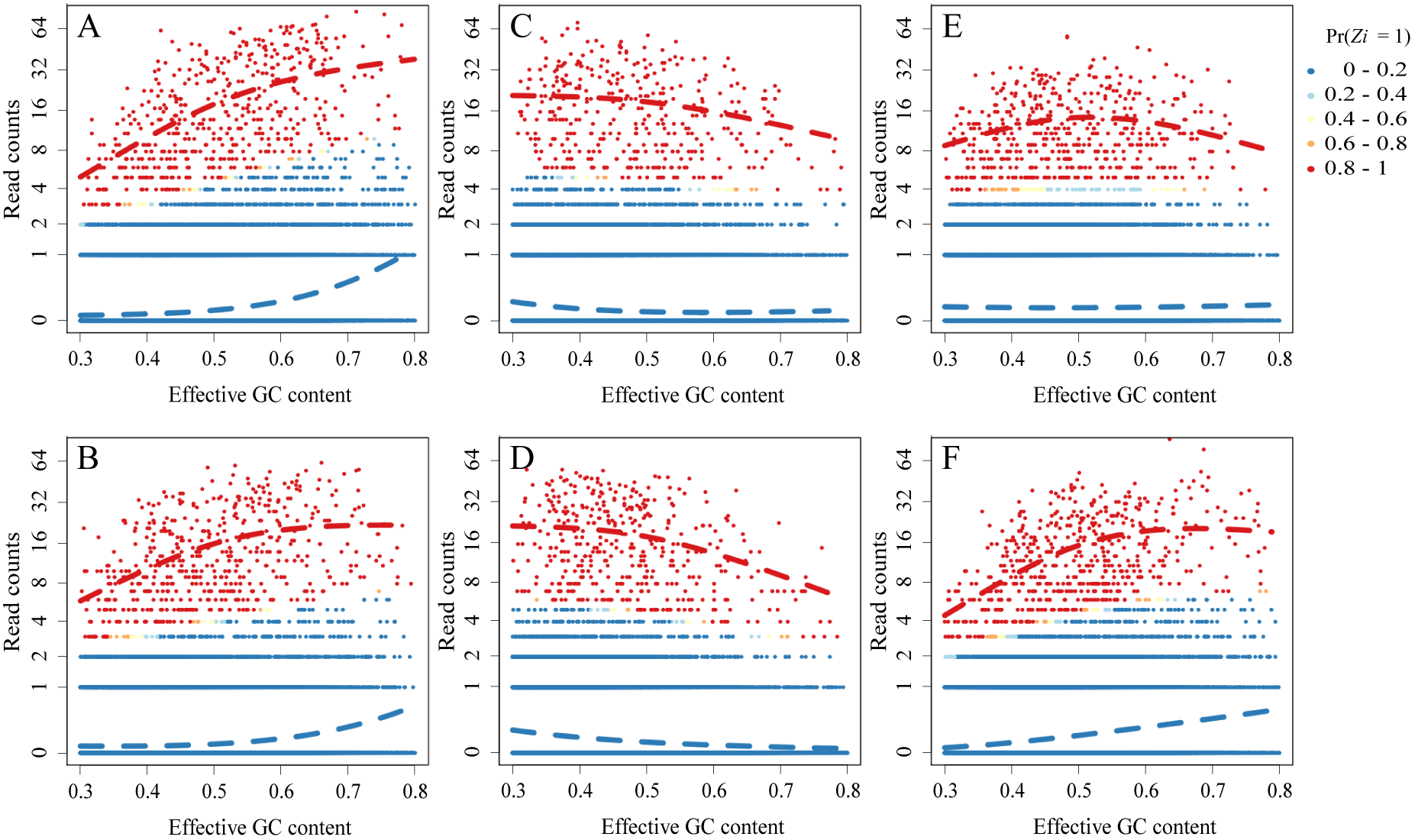
Visualization of the fitted generalized linear mixture model. A) We defined bins using estimated binding size and randomly selected a 5% of all genome-wide bins. We computed counts for these bins in the first replicated of the HUVEC cell line for laboratory UW. For these bins we fitted our model. The colors represent the probability of being background (blue) or signal (red). The GC-content bias smooth functions are plotted with dashed curves. B) As A) but for the second replicate for laboratory UW. C) As A) but for laboratory UTA. D) As C) but for the second replicate for laboratory UTA. E) As A) but for laboratory Broad. F) As E) but for the second replicate for laboratory Broad.

### Adjusting binding quantification for GC-bias reduces batch effects

We computed counts for each the regions reported as CTCF peaks in at least one cell line by ENCODE (Kundaje et al. 2015) for each of the GM12878, HeLa-S3, HepG2, HUVEC, K562 and NHEK samples. We constructed a matrix with these binding quantifications and performed principal component analysis (PCA) on the log-transformed values in this matrix. The first two principal components (PCs) do not separate by cell line as expected (Fig. 5A). Furthermore, the large variation seen within each cell line is largely explained by the different labs (Fig. 5A). We then adjusted the values in this matrix for GC-content using our model based approach and recomputed the PC_s_. The results were markedly improved (Fig. 5B) with the samples now clearly clustering by cell line and much of the batch effects removed. The improvement in specificity and batch effect removal was evident from plotting mean squared residuals summarizing across laboratory variability, computed within cell-line, before and after GC-content correction and noting a substantial reduction (Fig. 5C).

**Figure 5.**
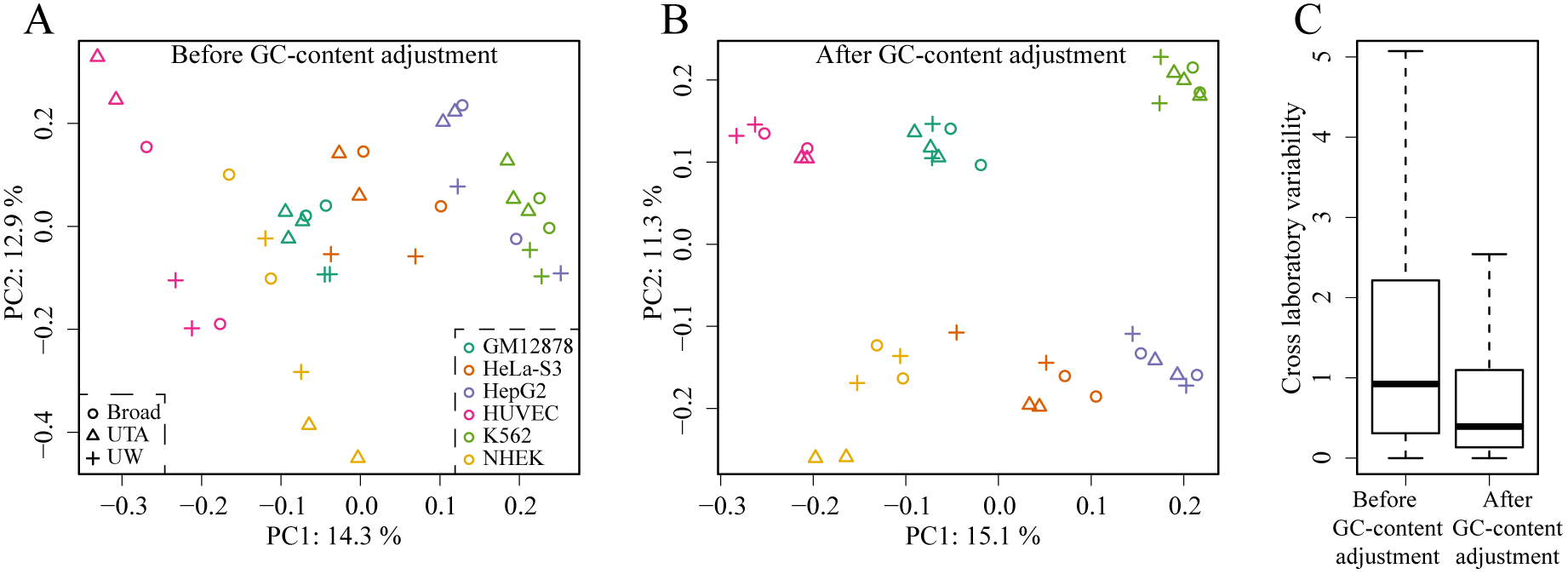
GC-bias correction reduces the impact of batch effects. A) For the regions reported as binding sites by ENCODE we computed counts for the GM12878, HeLa-S3, HepG2, HUVEC, K562 and NHEK. The first two principal components of this matrix are shown with color representing cell-line and different symbols used to represent lab. B) As A) but after performing the batch correction. C) Boxplots showing the within cell-line across laboratory variability before and after correction for the HUVEC cell line.

### Integrating GC-content adjustment into peak calling algorithms

Our model-based approach provides an adjustment value for any genomic bin. This implies that it can be integrated with peak algorithms that use bins as raw data (Ji et al. 2008; Kharchenko et al. 2008; Zhang et al. 2008; Rashid et al. 2011). Here we demonstrate the advantages of our approach by adapting the peak detection algorithm used by ENCODE, namely the ChIP-Seq processing pipeline (SPP) (Kharchenko et al. 2008). SPP starts by estimating the average half width of the binding protein, referred to here as *W* (see Methods Section for details). With this estimate in place the SPP algorithm then computes read counts for positive and negative strands separately for each genomic location *i*, denoted here with *Y_i, +_* and *Y_t, −_* respectively. The *Y_i, +_* represents positive strand counts in a region starting at *i* – *W* and ending at *i* and the *Y_t, −_* represents negative strand counts in a region starting at *i* and ending at *i* + *W*. As described by Pepke et al (Pepke et al. 2009), these counts should be large when a protein binds a region centered at *i*. To account for local background, SPP also computes counts in regions that should be associated with non-specific binding, denoted here with *B_i,+_* and *B_t,−_* respectively. The background level *B_i,+_* represents positive strand counts in a region starting at *i* and ending at *i* + *W* and the background level *B_t,−_* represnets negative strand counts in a region starting at *i* – *W* and ending at *i*. As described by Pepke et al (Pepke et al. 2009), there should be no counts in these regions when a protein binds a region centered at *i*. Then for each *i*, SPP defines the *enrichment score* as a geometric mean of the signal counts minus the average background signal 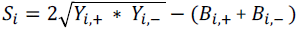. Note that this is geometric average of the signal minus the arithmetic average of background multiplied by 2. To find candidate peaks SPP then estimates of binding significance, and uses the local maxima of enrichment score to call peaks. To correct for GC we simply compute the effective GC-content of each bin and adjust *Y_i,+_*, *Y_i,−_*, *B_i,+_*, *B_i,−_* accordingly. We then used an approach similar SPP to quantify uncertainty for each candidate peak reported by our GC-corrected SPP algorithm (Methods and Supplemental Fig. 5). We compared these peaks to those obtained by the original SPP and found that our method (gcapc) resulted in substantial improvement in consistency (Fig. 6A). If, as done by ENCODE, we filter peaks using an IDR(Li et al. 2011) of 0.02, our algorithm reports improved results of 33,216, 35,234 and 30,089 peaks (Supplemental Fig. 2B) called for the three laboratories and now only 18.6% of regions reported by only one lab. More importantly the differences are no longer due to differences in GC-content (Fig. 2B). Note that the ENCODE pipeline is more complicated than running a peak caller and IDR (https://www.encodeproject.org/chip-seq/transcription_factor/). If we simply run SPP followed by IDR the improvements of our algorithm are even larger since this approach produced 29.5% regions reported by only one lab (Supplemental Fig. 2C)

Finally, to further assess the improvement provided by our approach we performed CTCF binding site enrichment analysis. Specifically, we used the most recent prediction of the human CTCF motif (Schmidt et al. 2012) to define a position weight matrix (PWM) score sequence of the same size as the motif (Wasserman and Sandelin 2004). Then we assigned a PWM score to each reported peak by selecting the maximum PWM within the region associated with the peak. Following (Wasserman and Sandelin 2004), we defined peaks with a maximum PWM score lower than 72% as a false positive. The gcapc had substantially less false positives than SPP (Fig. 6B). For example, if we examine the top 100 peaks across all six replicates, SPP results in a total of 17 false positives while gcapc has none.

**Figure 6.**
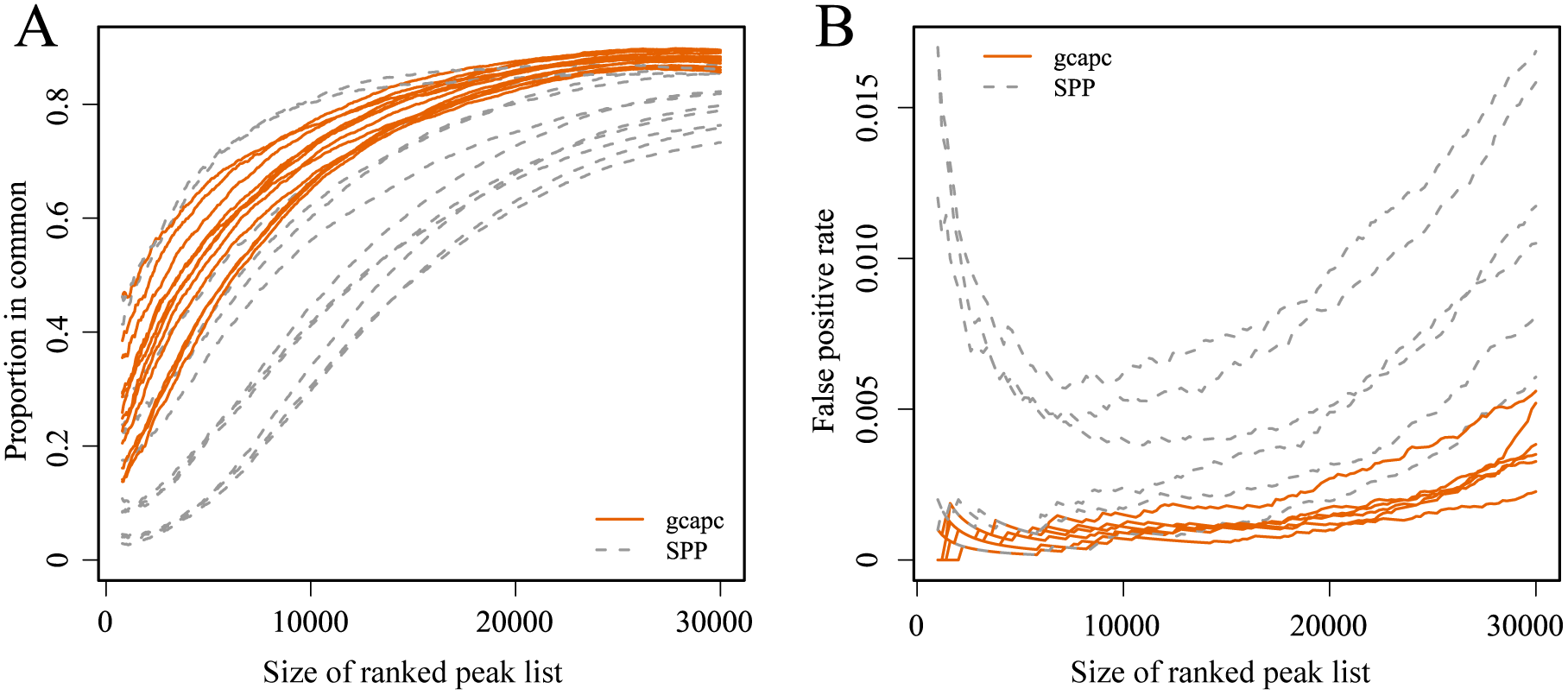
Improvements of peak calling consistency and TFBS enrichment with GC content adjustment. For the HUVEC cell-line we create lists of candidate peaks of varying. A) Correspondence at the top (CAT) plots. For each list size we compute the proportion of peaks in common reported by two different laboratories. We do this for each pairwise comparison and plot this percentage against the list size. Peak width is scaled to the same median size between gcapc and SPP for each sample. B) For each list size we compute PWM scores for each peak and define with a lower than 72% as false positives. We do this for each replicate and plot false positive rate against the list size and plot the number of false positives for each list sizes ranging from 1 to 30,000.

## Discussion

We have demonstrated how GC-content bias induces substantial variability into ChIP-Seq data and that this variability is large enough to result in different peaks being reported by different laboratories when studying the same cell-lines. We described how published GC-content adjustment methods are not directly applicable to ChIP-Seq data due to confounding between GC-content of regions and their biological relevance. We described *gcapc* (http://bioconductor.org/packages/gcapc/), a method that adjusts for GC-content bias in ChIP-Seq data using a mixed model, which permits independent adjustments of the signal and background signals and thus circumvents the confounding challenge, and can be incorporated into most current peak callers. Our method permits the GC-content bias correction for any predefined bin. We demonstrated the practical advantage of our approach by removing batch effects from binding quantifications in ENCODE data and by adapting the widely used SPP algorithm and showing substantial improvements in peak calling consistency across labs.

## Methods

### Data Acquisition and Preprocessing

We chose data for the transcription factor CTCF provided by ENCODE an example dataset because it includes a wide range of cell types and experiments performed by three different laboratories using the same protocol. The three laboratories were located at the Broad Institute (Broad), University of Texas at Austin (UTA) and University of Washington (UW). We focused on the GM12878, HeLa-S3, HepG2, HUVEC, K562 and NHEK cell lines because each of these was processed in replicates by each of the three laboratories. Raw sequencing reads were downloaded from ENCODE data portal (https://www.encodeproject.org/) using accession IDs documented in Supplemental Table 1.

The raw reads were aligned to human genome build hg19 with aligner BWA (Li and Durbin 2009). Reads from chromosome Y were ignored to avoid sex effects. Mapped reads with mapping score less than 30 were removed. Secondary alignments were also removed. Duplicate reads were thinned down to one read. For the purposes of quantifying binding in predefined region, we only considered the start position at the 5′ end.

### Estimating GC-content bias with mixed generalized linear model

Figure 1 and 4 clearly demonstrate the presence of two clusters. We assume the cluster characterized by low counts is related to non-specific binding and refer to it as the *background*. We assume that the cluster characterized by higher counts is related to the specific binding signals that constitute the peaks. The counts in both clusters show a strong non-linear dependence on GC-content and motivate the following mixed model. We assume that for any given position *i, Z_i_* = 1 if binding occurs at that position and 0 otherwise. We denote with *π* the probability that any given *Z_i_* = 1. We then assume that conditioned on the state *Z_i_*, the counts *Y_i_* follow a Poisson distribution with log (E[*Y_i_* | *Z_i_* = *a*, *X_i_* = *x_i_*]) = *μ_a_* + *f_a_*(*x_i_*) with *μ_a_* the mean count level for the positions, *x_i_* the effective GC content for position *i, f_a_* is a smooth function that we represent with a cubic spline. Note that *a* is indexing the two possible states, background or specific signal, which implies the GC-content effect is modeled differently for each state.

Because we start with millions of bins, to improve computational efficiency, we selected a random subset of bins representing 5% of the genome. We then estimated the parameters *π*, *μ*_0_, *μ*_1_ and the parameters used to represent the splines *f*_0_ and *f*_1_ using an EM algorithm. The GC-content effect used in the correction for binding quantification is simply

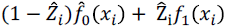

with *Ẑ_i_* = Pr (*Z_i_* = 1). To extend SPP we use the correction 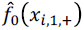 and 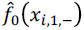 for the *Y_i_*,_+_ and *Y_i, −_* respectively where *x_i, 1, +_* and *x_i, 1, −_* are the effective GC-content in the positive and negative strand bins, described above, respectively. Similarly, we used the correction *f*_0_(*x_i, 0, +_*) and *f*_0_(*x_i, 0, −_*) for the *B_i, +_* and *B_i, −_* respectively. Note that we use *f*_0_ for both signal and background components because this summary is intended as a test statistic for which we define a null distribution assuming there is no signal. The use of *f*_1_ is therefore only used to fit the model and also for binding quantification of regions already determined to be potential peaks.

### Analysis of regions reported by ENCODE

For the analysis involving the binding regions reported by ENCODE (https://www.encodeproject.org/data/annotations/v2/) we did not need to run a peek-calling algorithm since regions were already provided. We used data from all the regions reported by ENOCDE as potential peaks. To perform a GC-content bias correction of binding quantification we assumed a binding width of 150bp and used flanking regions of width of 250bp. We fit the model described above and we can correct as described. PCA analysis was based on binding regions reported for GM12878, HeLa-S3, HepG2, HUVEC, K562 and NHEK cell lines. As an example, the across laboratory variability was computed in the HUVEC.

### Quantifying Uncertainty

We implement a method similar to SPP. Specifically, we compute the enrichment score *S_i_* for each region *i*. Then, for each of these regions, we permuted the start sites of all the reads falling the region and recomputed the enrichment scores, denoted here with 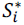. We used the 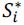 to form a null distribution and assign a p-value to each candidate peak (Supplemental Fig. 5). The user should treat the p-values obtained from this procedure, as well as SPP, with caution as they are based on several assumptions that are hard to test empirical. Furthermore, these uncertainty estimates do not account for the selection process. Permutation approaches such as those implemented by the bumphunter approach (Aryee et al. 2014) are the subject of future research. Regardless, we find this quantification useful for prioritizing peaks.

### Other improvements on SPP algorithm

Apart from the GC-content correction we adapted the SPP algorithm in three other ways, which we describe in detail here.

The SPP algorithm computes the enrichment score *S_i_* for every location *i* on the genome. To do this, SPP defines a window size *W* that is used to compute the read counts as described in the Results Section (Fig. 3). This window size is supposed to define the region that includes fragments resulting with a protein binding at location *i*. To define *W*, SPP uses the cross-correlation function between the fragment start sights from positive and negative strands. SPP uses an ad-hoc procedure picking *W* to be a number between the lag that maximizes the cross-correlation function and a use defined maximum window size that defaults to 500 basepairs. Instead, we make the assumption that the window size *W* should maximize the correlation of read counts between positive-strand windows and corresponding negative-strand windows. Using this criteria, the estimated window size in CTCF dataset are always much smaller than those estimated by SPP.

The second difference related to computational efficiency. While SPP computes *S_i_* and uncertainty estimate for every location on the genome. In our software we perform a filter that removes regions with small counts in all four relevant bins (*Y_i,+_*, *Y_i,−_*, *B_i,+_*, *B_t,−_*). This reduces the number of regions for which the uncertainty quantification is computed.

Finally, to report the center of the binding site, SPP searches for local maxima of enrichment scores *S_i_*. However, distributions of enrichment scores are not always symmetric around these local maxima, and we find that neighbored peaks regions sometimes represent the same binding site. To avoid reporting two maxima associated with the same binding site as two separate peaks, we merged any two neighboring peaks that are within the estimated binding width from each other. Uncertainty is quantified only for merged peaks.

### Software and source code availability

The method described in this manuscript is available as an R/Bioconductor package (http://bioconductor.org/packages/gcapc/). The source code for the main results is documented here (https://github.com/tengmx/gcapc_manuscript).

## Acknowledgments

We thank Anshul Kundaje for the suggestion on transcription factor binding sites analysis.

## References

Aird D, Ross MG, Chen WS, Danielsson M, Fennell T, Russ C, Jaffe DB, Nusbaum C, Gnirke A. 2011. Analyzing and minimizing PCR amplification bias in Illumina sequencing libraries. Genome Biol 12: R18.

Alkan C, Kidd JM, Marques-Bonet T, Aksay G, Antonacci F, Hormozdiari F, Kitzman JO, Baker C, Malig M, Mutlu O et al. 2009. Personalized copy number and segmental duplication maps using next-generation sequencing. Nat Genet 41:1061–1067.

Aryee MJ, Jaffe AE, Corrada-Bravo H, Ladd-Acosta C, Feinberg AP, Hansen KD, Irizarry RA. 2014. Minfi: a flexible and comprehensive Bioconductor package for the analysis of Infinium DNA methylation microarrays. Bioinformatics 30:1363–1369.

Benjamini Y, Speed TP. 2012. Summarizing and correcting the GC content bias in high-throughput sequencing. Nucleic Acids Res 40: e72.

Celniker SE, Dillon LA, Gerstein MB, Gunsalus KC, Henikoff S, Karpen GH, Kellis M, Lai EC, Lieb JD, MacAlpine DM et al. 2009. Unlocking the secrets of the genome. Nature 459: 927–930.

Cheung MS, Down TA, Latorre I, Ahringer J. 2011. Systematic bias in high-throughput sequencing data and its correction by BEADS. Nucleic Acids Res 39: e103.

Dohm JC, Lottaz C, Borodina T, Himmelbauer H. 2008. Substantial biases in ultra-short read data sets from high-throughput DNA sequencing. Nucleic Acids Res 36: e105.

Dunham I, Kundaje A, Aldred SF, Collins PJ, Davis CA, Doyle F, Epstein CB, Frietze S, Harrow J, Kaul R et al. 2012. An integrated encyclopedia of DNA elements in the human genome. Nature 489: 57–74.

Hansen KD, Irizarry RA, Wu Z. 2012. Removing technical variability in RNA-seq data using conditional quantile normalization. Biostatistics 13: 204–216.

Ji H, Jiang H, Ma W, Johnson DS, Myers RM, Wong WH. 2008. An integrated software system for analyzing ChIP-chip and ChIP-seq data. Nat Biotechnol 26:1293–1300.

Jothi R, Cuddapah S, Barski A, Cui K, Zhao K. 2008. Genome-wide identification of in vivo protein-DNA binding sites from ChIP-Seq data. Nucleic Acids Res 36: 5221–5231.

Kharchenko PV, Tolstorukov MY, Park PJ. 2008. Design and analysis of ChIP-seq experiments for DNA-binding proteins. Nat Biotechnol 26:1351–1359.

Kundaje A, Meuleman W, Ernst J, Bilenky M, Yen A, Heravi-Moussavi A, Kheradpour P, Zhang Z, Wang J, Ziller MJ et al. 2015. Integrative analysis of 111 reference human epigenomes. Nature 518: 317–330.

Landt SG, Marinov GK, Kundaje A, Kheradpour P, Pauli F, Batzoglou S, Bernstein BE, Bickel P, Brown JB, Cayting P et al. 2012. ChIP-seq guidelines and practices of the ENCODE and modENCODE consortia. Genome Res 22:1813–1831.

Li H, Durbin R. 2009. Fast and accurate short read alignment with Burrows-Wheeler transform. Bioinformatics 25:1754–1760.

Li Q, Brown JB, Huang H, Bickel PJ. 2011. Measuring reproducibility of high-throughput experiments. doi:10.1214/11-AOAS466:1752–1779.

Love MI, Hogenesch JB, Irizarry RA. 2016. Modeling of RNA-seq fragment sequence bias reduces systematic errors in transcript abundance estimation. Nat Biotechnol doi:10.1038/nbt.3682.

Pepke S, Wold B, Mortazavi A. 2009. Computation for ChIP-seq and RNA-seq studies. Nat Methods 6: S22–32.

Rashid NU, Giresi PG, Ibrahim JG, Sun W, Lieb JD. 2011. ZINBA integrates local covariates with DNA-seq data to identify broad and narrow regions of enrichment, even within amplified genomic regions. Genome Biol 12: R67.

Ross MG, Russ C, Costello M, Hollinger A, Lennon NJ, Hegarty R, Nusbaum C, Jaffe DB. 2013. Characterizing and measuring bias in sequence data. Genome Biol 14: R51.

Rozowsky J, Euskirchen G, Auerbach RK, Zhang ZD, Gibson T, Bjornson R, Carriero N, Snyder M, Gerstein MB. 2009. PeakSeq enables systematic scoring of ChIP-seq experiments relative to controls. Nat Biotechnol 27: 66–75.

Schmidt D, Schwalie PC, Wilson MD, Ballester B, Goncalves A, Kutter C, Brown GD, Marshall A, Flicek P, Odom DT. 2012. Waves of retrotransposon expansion remodel genome organization and CTCF binding in multiple mammalian lineages. Cell 148: 335–348.

Valouev A, Johnson DS, Sundquist A, Medina C, Anton E, Batzoglou S, Myers RM, Sidow A. 2008. Genome-wide analysis of transcription factor binding sites based on ChIP-Seq data. Nat Methods 5: 829–834.

Wasserman WW, Sandelin A. 2004. Applied bioinformatics for the identification of regulatory elements. Nat Rev Genet 5: 276–287.

Zhang Y, Liu T, Meyer CA, Eeckhoute J, Johnson DS, Bernstein BE, Nusbaum C, Myers RM, Brown M, Li W et al. 2008. Model-based analysis of ChIP-Seq (MACS). Genome Biol 9: R137.

